# The J domain of sacsin disrupts intermediate filament assembly

**DOI:** 10.1101/2021.07.14.452331

**Authors:** Afrooz Dabbaghizadeh, Alexandre Paré, Zacharie Cheng-Boivin, Robin Dagher, Sandra Minotti, Marie-Josée Dicaire, Brais Bernard, Jason C. Young, Heather D. Durham, Benoit J. Gentil

## Abstract

Autosomal Recessive Spastic Ataxia of the Charlevoix Saguenay (ARSACS), is caused by loss of function mutations in the *SACS* gene, which encodes sacsin, a giant protein of 520 kDa. A key feature of the absence of sacsin in cells is the formation of abnormal bundles of intermediate filaments (IF) including neurofilaments (NF) in neurons and vimentin IF in fibroblasts, suggesting a role of sacsin in IF homeostasis. Sacsin contains a J domain (SacsJ) homologous to Hsp40, that can interact with Hsp70 chaperones. The SacsJ domain resolved NF bundles in cultured *Sacs*^-/-^ neurons, however, its mechanism is still unclear. Here, we focused on the role of SacsJ in NF assembly. We report that the SacsJ domain directly interacts with NF proteins *in vitro* to disassemble NFL filaments, and to inhibit their initial assembly, in the absence of Hsp70. We generated a cell-penetrating peptide derived from this domain, SacsJ-myc-TAT, which was efficient in disassembling both vimentin IF and NF in cultured fibroblasts and *Sacs*^+/+^ motor neurons as well as NF bundles in cultured *Sacs*^-/-^ motor neurons. Whereas a normal NF network was restored in *Sacs*^-/-^ neurons treated with the SacsJ peptide, there was some loss of IF networks in *Sacs*^+/+^ fibroblasts or neurons. These results suggest that SacsJ is a key regulator of NF and IF networks in cells, with implications for its therapeutic use.

## Introduction

Autosomal Recessive Spastic Ataxia of the Charlevoix Saguenay (ARSACS), the second most common recessive ataxia worldwide, is a childhood-onset neurodegenerative disease caused by over 200 different mutations in the *SACS* gene. *SACS* encodes the giant 520 kDa, multi-domain protein, sacsin [1, 2]. Although ubiquitously expressed, sacsin is a cytoplasmic protein enriched in neurons [3]. ARSACS patients exhibit a clinical triad of ataxic gait, pyramidal spasticity, and peripheral neuropathy [4–9]. Sacsin knockout (*Sacs*^-/-^) mice exhibit a similar phenotype, supporting a loss of function mechanism [10].

The domain structure of sacsin, characterised based on sequence homology using *in silico* analysis [11], suggests functions in protein chaperoning and quality control. Sacsin contains, starting at the N-terminus, an ubiquitin-like domain (UBL), which can bind to the proteasome 20S α subunit [12]; three large sacsin repeat regions (SRR), termed SIRPT1, SIRPT2 and SIRPT3, which share homology with the chaperone, Hsp90; an XPC-binding domain; and then at the C-terminus a J domain (SacsJ) homologous to Hsp40, immediately followed by a single higher eukaryote and prokaryote nucleotide-binding domain (HEPN) believed to promote sacsin dimerization [11, 13-15].

Evidence, including homology to heat shock proteins, further points to chaperone and quality control roles of various sacsin domains. The XPC-binding domain of another protein, HR23A, binds to the ubiquitin ligase Ube3A, suggesting that sacsin might be ubiquitinylated by Ube3A [16]. Sacsin’s N-terminus including SIRPT1 exhibited chaperoning activity in a luciferase folding assay, cooperatively with Hsp40 [12]. The sacsin J domain can bind Hsp70 chaperone family proteins and resolved aggregates of ataxin1 variants when overexpressed, which suggests that SacsJ acts as a cochaperone [13]. Autophagy regulation is defective in SHSY5Y cells after knockout of the *SACS* gene and in fibroblasts derived from ARSACS patients [17], and autophagic flux was increased in patient-derived fibroblasts although the mechanism by which sacsin is involved in autophagy is not known [18]. In fact, molecular chaperones, such as sacsin, associated with folding/unfolding and assembly/disassembly of protein complexes have been considered as important players of selective autophagy [19].

A key feature of lack of sacsin is abnormal bundling of intermediate filaments (IF), including neurofilaments (NF), the neuron-specific intermediate filaments, in *Sacs*^-/-^ mice and cultured neurons and vimentin IF in cultured fibroblasts derived from patients’ skin biopsies [10, 18]. Mitochondrial elongation is also a characteristic of cells lacking sacsin [20, 21], although we showed mitochondrial abnormalities to be a later event than NF bundling in cultured *Sacs*^-/-^ motor neurons [10] and mitochondrial morphology in general to be dependent on a normal NF network [22]. We obtained direct evidence of sacsin’s function as a chaperone in IF assembly and homeostasis in cultured *Sacs*^-/-^ motor neurons and in patient-derived fibroblasts by ectopic expression of sacsin domains [23]. Subtle differences in the NF network were observed according to which of the sacsin domains was expressed: the UBL domain decreased NF bundles and the amount of NF proteins, without affecting NF assembly *per se*; SIRPT domains provided a scaffold for assembly of NF proteins into long filaments; expression of the SacsJ domain prevented assembly of NF, and expression of the HEPN domain re-localized filaments to the periphery of the cell forming a cage-like structure [23].

IF are dynamic structures that undergo assembly/disassembly and annealing/severing. IF assembly is an organised stepwise process leading to the formation of 10 nm thick non-polar filaments [24]. IF proteins assemble into dimers through interaction of their coiled-coil domains, which then polymerise into tetramers. Eight tetramers assemble laterally to form a cylindrical Unit-Length Filaments (ULF) (~60 nm long), which anneal with other ULF to form an immature filament [25]. Radial compaction of the filament is the last step to form mature 10 nm diameter filaments, which can be solubilized in 8M urea [26, 27]. Although post-translational modifications play a regulatory role in IF assembly [28], so do a subset of chaperone proteins. For example, the small heat shock proteins HspB1/Hsp27 and HspB5/αB-crystallin decreased the rate of NFL polymerization and affected NFL transition from tetramers to filaments and their bundling in vitro [29]. HSPB5/αB-crystallin prevents IF assembly and network formation [30–32], HSPB1/Hsp27 modifies the IF assembly dynamics, preventing extensive bundling of NF and maintaining an ordered NF assembly [33]. Finally, HSPA1/Hsp70 can properly fold Charcot-Marie-Tooth type 2E-causing NFL variants into a normal filamentous network [22], but few co-chaperones like DNAJB6/Mrj have IF proteins as clients [34].

We previously reported that expression of the SacsJ domain resolved NF bundles in motor neurons cultured from *Sacs*^-/-^ mice [23]. Compared to the UBL and SIRPT1 domains, SacsJ was the most effective in these neurons. In fibroblasts, SacsJ also promoted disassembly of the vimentin IF network [23]. Although SacsJ can act as a co-chaperone of Hsp70, it also has some chaperone activity of its own [13]. Therefore, SacsJ may be acting directly on filaments without the involvement of Hsp70. This activity may be as a general chaperone, or a chaperone with specific effects on NF.

These questions were addressed in the present study. We tested whether a peptide corresponding to the SacsJ domain (aa 4316-4420) regulates IF assembly directly or indirectly, and whether it has general chaperoning function on its own. In an in vitro NF assembly system, SacsJ both promoted dissolution of NF assembled from recombinant human NFL and inhibited their assembly, pointing to a direct role for this domain of sacsin in NF dynamics. We developed a cell-penetrating peptide to further examine SacsJ’s effect on IF in vivo. The cell-penetrating peptide SacsJ-myc-TAT significantly resolved NF bundles in *Sacs*^-/-^ motor neurons, but disassembled the IF network in *Sacs*^+/+^ fibroblasts or cultured motor neurons. These data suggest that SacsJ is a potent regulator of IF networks, with implications for its possible therapeutic uses.

## Materials and Methods

### Cloning, protein production and purification

The SacsJ cDNA (corresponding to residues 4316-4420) inserted in frame with GST into the pGEX6 vector was amplified from the mouse pEGFP-sacsin full length (OriGene Technologies, Rockville, MD, USA). For delivery into cells and tissues, the pGEX-SacsJ or GST were tagged on the C-terminus with a myc epitope to identify the peptide and with the TAT-derived cell-penetrating peptide sequence (YGRKKRRQRRR) [35], which has been shown to confer efficient neuronal delivery and blood-brain barrier penetration [36]. All cloning was subcontracted to NorClone (London, Ontario, Canada [NorClone - Gene Cloning Supplier](http://norclone.com/)). Human NFL cloned into the pET23b expression vector was a gift of Dr. Walter Mushynski (retired, McGill University).

For recombinant protein production, plasmids carrying the SacsJ-myc-TAT or GST-myc-TAT cDNA were transformed into the Escherichia coli BL21 (DE3) pLysS. Bacterial cultures were grown overnight at 37 °C and then diluted at 1:100 (v/v) into 1L Luria–Bertani medium containing 100μg/ml ampicillin. Bacteria were grown at 37 °C with vigorous shaking at 225 rpm until they reach an OD600 nm=0.6, recombinant protein expression was then induced using 1 mM isopropyl-1-thio-β-D galactopyranoside (IPTG) at 30 °C for 4 hours.

Bacteria were lysed in PBS buffer (137 mM NaCl, 2.7 mM KCl, 10 mM Na_2_HPO_4_, 2 mM KH_2_PO_4_ pH 7.4), supplemented with 1 mM phenylmethylsulfonyl fluoride (PMSF), 80 units of DNase,100 μg/ml lysozyme and Triton 0.1 %. After sonication, bacterial debris was pelleted by centrifugation. The supernatant was then filtered through a 0.2 μm filter and chromatography affinity purification of GST-myc-TAT or GST-SacsJ-myc-TAT using glutathione resin was carried out according to the manufacturer instructions (GE-Healthcare).

Production of recombinant human NFL was induced in the presence of 0.4 mM IPTG for 4h at 37° C according to Leung and Liem [37]. Briefly, after bacteria lysis, NFL was recovered in inclusion bodies. The insoluble protein pellet containing NFL was solubilised in Buffer 1 (8 M urea, 10 mM sodium phosphate buffer, pH 7.4, 0.1 % (v/v) 2-mercaptoethanol and 1x proteasome inhibitors from Sigma) and centrifuged. The supernatant was filtered through a 0.45-μm syringe filters and solubilized NFL was then purified from the supernatant by affinity chromatography using Bio-gel^®^ HT hydroxyapatite (130-0150, Bio-Rad) and following the protocol of Leung and Liem [37]. The column was then washed with Buffer 2 (8 M urea, 100 mM sodium phosphate buffer, pH 7, 0.1 % (v/v) 2-mercaptoethanol and 1x proteases inhibitors) and NFL was eluted in Buffer 3 (8 M urea, 300 mM sodium phosphate buffer, pH 7, 0.1 % (v/v) 2-mercaptoethanol and 1x proteases inhibitors).

### *In vitro* assembly of NFL

The assembly of NFL followed the procedure of Leung and Liem [37]. Urea was gradually removed by dialysis against the assembly buffer (50 mM MES, 0.5 mM EGTA, 0.175 M NaCl, 1 mM DTT, 0.4 mM PMSF, pH 6.8) containing 4M urea, 2M urea or no urea using a D-Tube Dialyzer MWCO 6-8 kDa (Millipore Sigma). NFL assembly (0.24 μg/μl total) was achieved by overnight dialysis at 4 °C. Filament assembly was assessed by the formation of filamentous structures observed by electron microscopy and by biochemical analysis using the insoluble properties of assembled NFL. After overnight dialysis, the solution containing assembled NFL was incubated with or without GST-SacsJ for 12h and then centrifuged at 100,000g for 30 min at 20 °C to separate the soluble and insoluble fractions, containing respectively soluble NFL and dimers or filamentous NFL. The soluble and insoluble fractions were then analysed by SDS-PAGE using Coomassie Blue staining.

### Negative staining and Electron Microscopy

Electron microscopy was performed at the Facility for Electron Microscopy Research (FEMR) at McGill University. 10 μl of protein samples were deposited and adsorbed for 1 min onto carbon-coated grids that were glow discharged before the application of proteins. Grids were Glow discharged for 1-2 min in a vacuum evaporator (Edwards Vacuum Carbon Coater E306) before adding the protein samples to improve quality of sample analysis. Assembled NFL as well as SacsJ were diluted to 0.24 μg/μl and processed for imaging by TEM. Excess of protein was removed by wicking the edge of the grid on a piece of Whatman paper. The samples were then negatively stained for 1 min using 10 μl of 2% (w/v) uranyl acetate and examined with a Cryo-TEM (FEI Tecnai G2 Spirit Twin) (Thermo Fisher Scientific) using an accelerated voltage of 120 KV. Images were acquired with a CCD camera (Gatan Ultrascan 4000 4k x 4k CCD Camera System Model 895). The diameters of the assembled filaments were measured on enlarged TEM micrographs using ImageJ software (National Institute of Health, Bethesda, MD, USA; https://imagej.nih.gov/ij/).

### Cell culture

Human skin fibroblasts were from the CellBank Repository for Mutant Human Cell Strains (McGill University Health Complex, Montreal, QC, Canada). Immortalised fibroblasts were cultured in Dulbecco’s Modified Essential Medium with 10% fetal bovine serum (FBS). Cultures were treated were treated with GST-myc-TAT or GST-SacsJ-myc-TAT to assess the time-dependence (30 min to 24h) and dose-dependence (0-5 μM) of the peptides effects on the vimentin IF network, visualized by immunolabeling with antibody to vimentin (clone V9, MA5-11883 Thermofisher). Peptides were identified by immunolabeling using an anti-myc antibody (C3956, Sigma-Aldrich). Cy2 or Cy3 conjugated donkey secondary antibodies against mouse or rabbit IgG were from Jackson ImmunoResearch (1/300).

Primary cultures of dissociated spinal cord-dorsal root ganglia (DRG) were prepared from E13 Sacs^-/-^ mice (C57Bl6 background) and wild type (*Sacs*^+/+^) of the same background. Generation and characterization of the Sacs^-/-^ mice were as previously described [10]. Cells were plated on glass coverslips (Fisher, ON, Toronto, Canada) coated with poly-D-lysine (P7280, Sigma-Aldrich) and Matrigel^®^ (CACB354234, VWR, Town of Mount Royal, QC, Canada) and maintained in Eagle’s Minimum Essential Medium enriched with 5 g/l glucose and supplemented with 3% horse serum, and other growth factors as previously described [38]. Cultures were used in experiments 6 weeks following plating to allow neuronal maturation and appearance of NF bundles in more than 80% of *Sacs*^-/-^ motor neurons. NF were visualized by immunolabeling with anti-NFL (clone NR4, N5139, Sigma-Aldrich). NF bundles were defined as previously by a continuous compact filament bundle crossing the cell body from dendrite to dendrite, distinguished from the normal finer NF network [23].

### In vitro chaperone assay

Chaperone activity of GST-SacsJ was determined in a heat-induced aggregation assay, measuring its ability to prevent heat denaturation of two protein substrates: pig citrate synthase (CS) (0.2 μM; MW 51 kDa; Sigma Aldrich), and catalase purified from bovine liver (2 μM; MW 250 kDa; Sigma Aldrich). Time duration for this assay was 90 min for CS and 50 min for catalase. Absorbance was measured every 10 min at 320 nm or 340 nm for CS or catalase, respectively. The substrates were heat-denatured at 45 °C for CS and at 60 °C for catalase in the absence or presence of SacsJ equal to 0.1-0.6 μM for CS and 1-6 μM for catalase. All assays were performed in 96-well clear-bottom plates (Sarstedt) using a spectrophotometer with thermostated cells (Varian Cary 100, Montreal, Quebec, Canada) in HEPES buffer (50 mM HEPES-KOH, pH 7.5) for CS and Sodium Phosphate buffer (100 mM, pH 7.5) for catalase [39]. Data are representative of three independent assays and are expressed as the mean ± SD.

### Statistics

Quantitative experiments are presented as mean ± SD. Length and number of NFL filaments were measured on TEM images taken from at least two separate dialysis batches. The proportion of motor neurons or fibroblasts carrying a dismantled, regular or bundled IF network was quantified in at least 3 coverslips per condition, with a minimum count of 30 cells per coverslip. P values < 0.05 were considered statistically significant. Statistical analysis was performed by using one-way ANOVA followed by a Tuckey HSD post hoc-analysis.

## Results

### SacsJ disassembles NFL filaments in vitro

Ectopic expression of the SacsJ domain in *Sacs*^-/-^ motor neurons in culture resolved NF bundles [23]. To determine if the SacsJ domain can alter NF properties directly, i.e., was its effect in cultured neurons due to a direct interaction with NF, we used an in vitro NF assembly assay. In vitro assembly of IF has been extensively studied and examination of the negatively stained filaments by transmission electron microscopy (TEM) is a standard method of characterization [37]. Purified recombinant human NFL incubated in assembly buffer formed filaments and some shorter filaments of 60 nm long that could correspond to UFL (Fig. 1A); GST-SacsJ alone did not form filamentous structures following incubationin NFL assembly buffer (Fig. 1B). After being assembled, NFL filaments were incubated with GST-SacsJ or GST as control. GST-SacsJ dismantled the pre-existing NFL filaments, as shown by the decreased average length and number of filaments observed by TEM (Fig. 1C and Fig. 1E).

**Figure 1:**
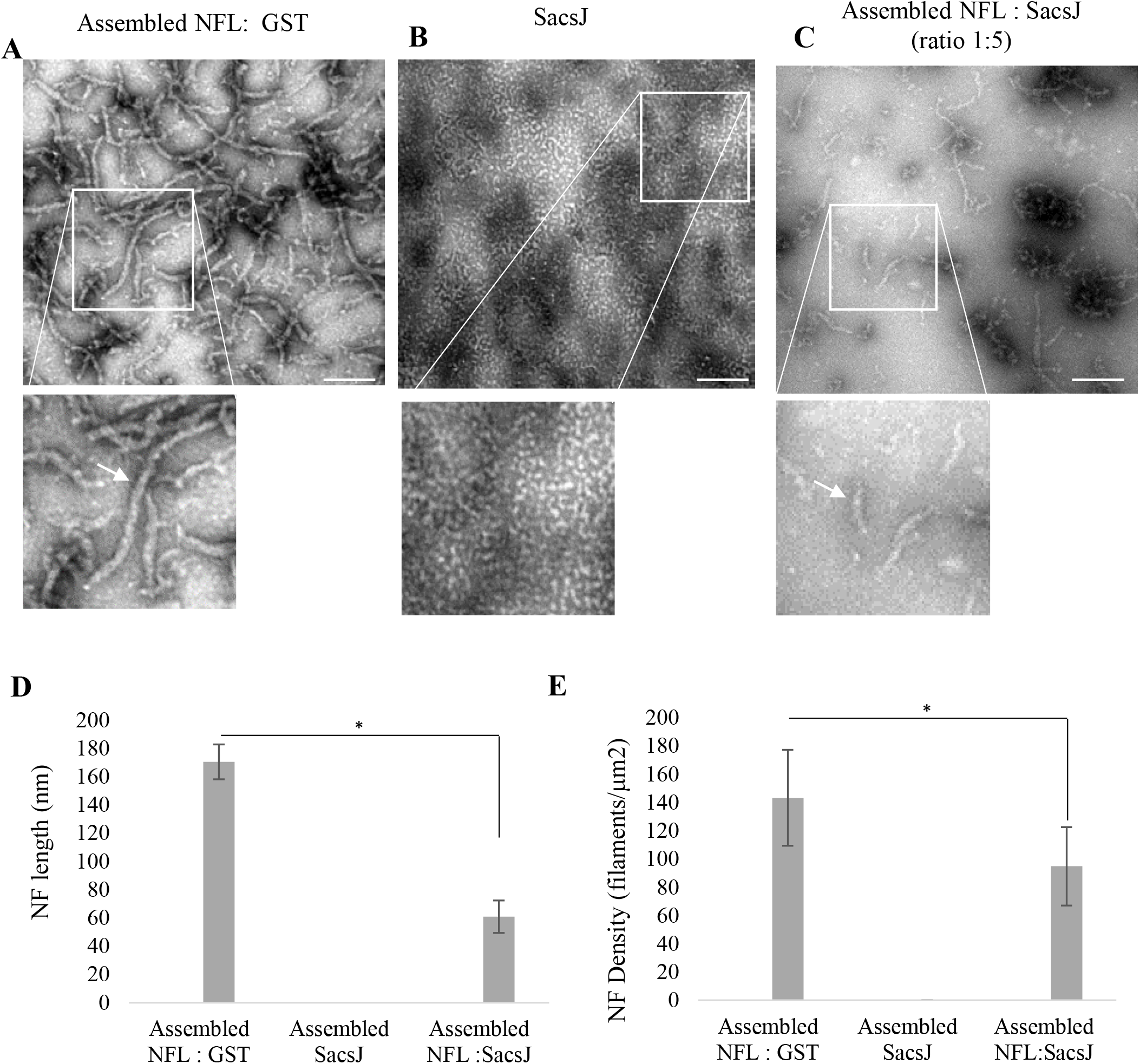
SacsJ dissembled NFL filaments in vitro. Shown are representative TEM images of filaments assembled from purified NFL (A). Assembled NFL filaments were incubated with or without SacsJ for 1h at 37 °C and observed by TEM. Representative TEM images of filamentous NFL (A), SacsJ alone (B) or filamentous NFL co-incubated with SacsJ at a molar ratio of 1:5 (C). Inserts show enlargements of TEM images focussed on NFL filaments (arrow). Scale Bar: 40nm. Quantitation of NFL average length (D) and filament density with or without incubation with SacsJ shows the significant decrease in NFL length and density (filaments/μm^2^). *p< 0.05 *vs* NFL alone, one-way ANOVA, HSD Tuckey post hoc analysis (n=3)

### SacsJ prevents in vitro assembly of NFL into filaments

To determine if SacsJ also prevents NF assembly, GST-SacsJ or GST were co-incubated with purified recombinant NFL in the NF assembly assay (Fig. 2). Incubation of NFL with GST-SacsJ at a ratio of 1:1 (NFL:SacsJ) reduced the assembly of NFL into filaments (Fig. 2B) as shown by the decrease of the average length of filamentous structures (Fig. 2F). Instead, NFL formed large insoluble, amorphous aggregates consistent with the structure and assembly of NFL being disrupted (Fig. 2C). Increased concentrations of GST-SacsJ at ratios of 5:1 and 10:1 totally prevented the formation of NFL filaments (Fig. 2C-E). The average length of the NFL filaments was under 60 nm when GST-SacsJ was incubated with NFL during in vitro assembly, suggesting that the SacsJ domain prevented the assembly of NFL into UFL (Fig. 2F).

**Figure 2:**
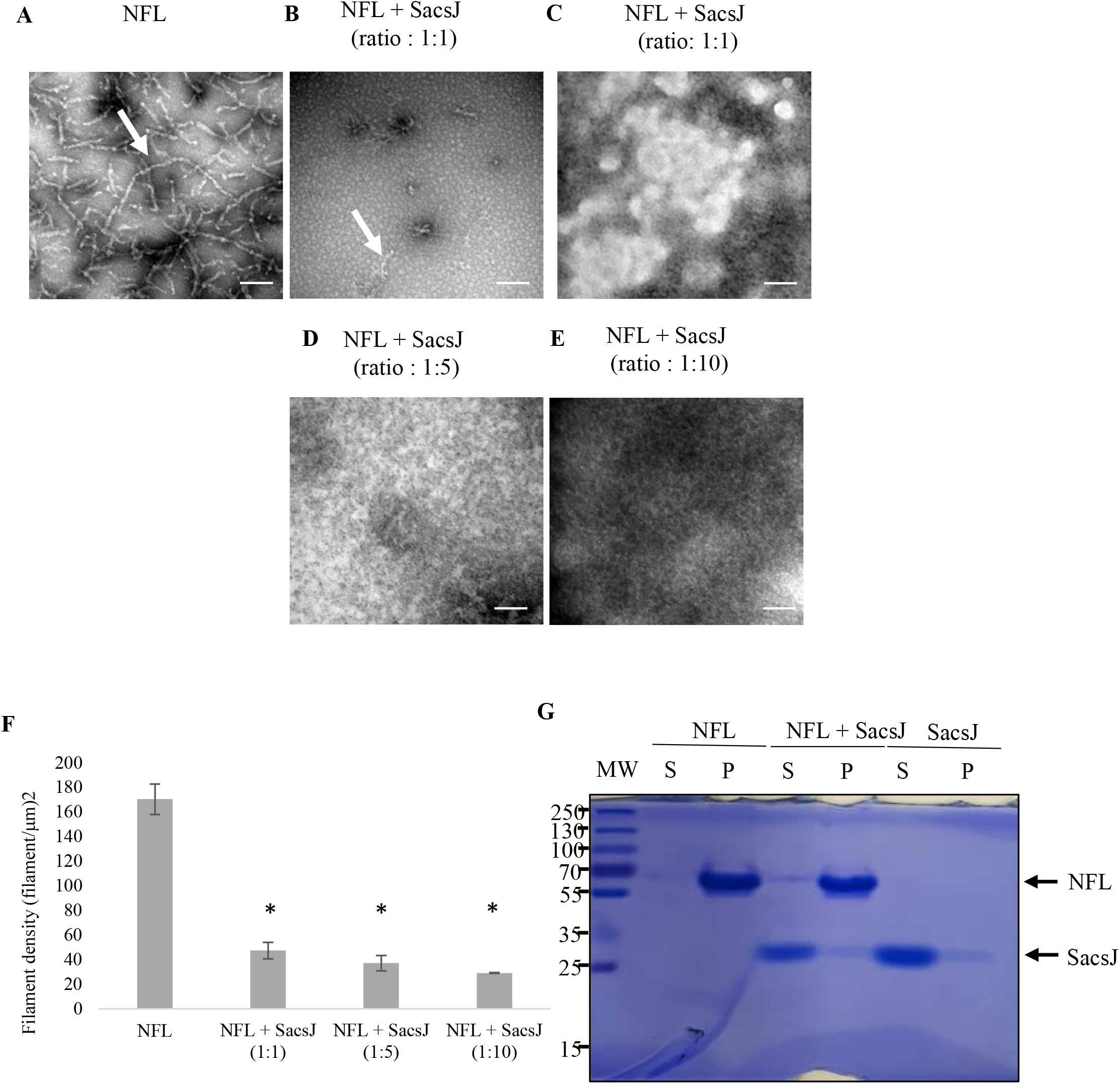
Effects of SacsJ on NFL assembly in vitro. A-D) TEM of negatively stained preparations of assembled NFL dialysed alone (A) or in the presence of different molar ratios of NFL to GST-SacsJ: 1:1 (B-C), 1:5 (D), and 1:10 (E). Scale bars: 100 nm. SacsJ prevented formation of NFL filaments (arrow). F) Quantitation of NFL filament density SacsJ shows significant reduction (filaments/μm^2^) by incubation with GST-SacsJ. *p< 0.05 *vs* NFL alone, one-way ANOVA (n=3). G) SDS-PAGE of a sedimentation assay of NFL with or without SacsJ in vitro after assembly at 4 °C overnight at ratio of 1:10. The supernatant (S) and pellet (P) fractions were analyzed by SDS-PAGE followed by Coomassie Blue staining. NFL and SacsJ bands are indicated by arrows and molecular weight is on the left.

Interestingly, the small heat shock proteins HSPB1/Hsp27 or HSPB5/αB-crystallin are known to have such an effect on in vitro assembly of IF, releasing small dimers and tetramers in the soluble fraction of a sedimentation assay while filamentous IF were found in the pellet [29, 31]. Therefore, the ability of SacsJ to change NFL sedimentation was assessed. After overnight incubation of NFL in assembly buffer (MES buffer) in vitro, with or without GST-SacsJ, supernatant and pellet fractions were separated by centrifugation and analysed by SDS-PAGE; gels were stained with Coomassie blue to visualize protein bands (Fig. 2G). Alone, NFL was mainly present in the pellet fraction (Pellet) and SacsJ in the supernatant fraction (Soluble), as expected from data presented in Fig. 1. Co-incubation of NFL with GST-SacsJ did not change the fraction in which NFL or SacsJ distributed; aggregates of NFL were retained in the pellet fraction, even though filament formation was inhibited, and SacsJ was retained in the supernatant. These data suggest that the interaction between SacsJ and NFL is transient because SacsJ did not co-sediment with NFL or switch fraction, which would indicate the formation of a stable complex like was formed between HSPB1 and NFL [29].

### The SacsJ domain lacks general chaperoning activity in vitro

Since the SacsJ domain alone altered NFL assembly in vitro, we tested whether it could function as a protein chaperone more generally. This was tested using an in vitro chaperoning assay based on heat-induced denaturation of catalase or citrate synthase (CS) [40]. Following heat-exposure, substrates such as CS and catalase unfold and form aggregates, measured by increased absorbance at 320 nm or 340 nm, respectively, which can be prevented by chaperone activity to prevent aggregation. Purified CS and catalase denature at different temperatures, 45 °C and 60 ᵒ°C within 90 or 60 minutes, respectively. In order to assess if SacsJ can prevent aggregation of substrates, GST (Fig. 3 C,D) or GST-SacsJ (Fig. 3 E,F) were mixed with catalase or CS at different molar ratios in the heat denaturation assay. At the molar ratio of 1:1 SacsJ:catalase, GST-SacsJ did not reduce the aggregation of catalase (Fig. 3E). Rather, increasing concentrations of GST-SacsJ exacerbated the aggregation of catalase, suggesting that SacsJ either aggregates itself or promotes unfolding and aggregation of the substrate. When tested alone in the assay, GST-SacsJ aggregated to some extent in a concentration-dependent fashion, but far below levels achieved by catalase (Fig. 3B); i.e., the increased absorbance obtained by combining catalase with SacsJ was much greater than the sum of absorbance measured individually. With CS as the substrate, SacsJ reduced its aggregation at a molar ratio of 1:2 SacsJ:CS (Fig. 3F); however, similar to experiments with catalase, the higher concentrations of GST-SacsJ increased CS aggregation (molar ratios 2:1 and 3:1). All together, our data confirm that SacsJ alone lacks significant inherent, general chaperoning activity. This suggests that the SacsJ effect on NFL formation may be due to specific disruption of NFL structure.

**Figure 3:**
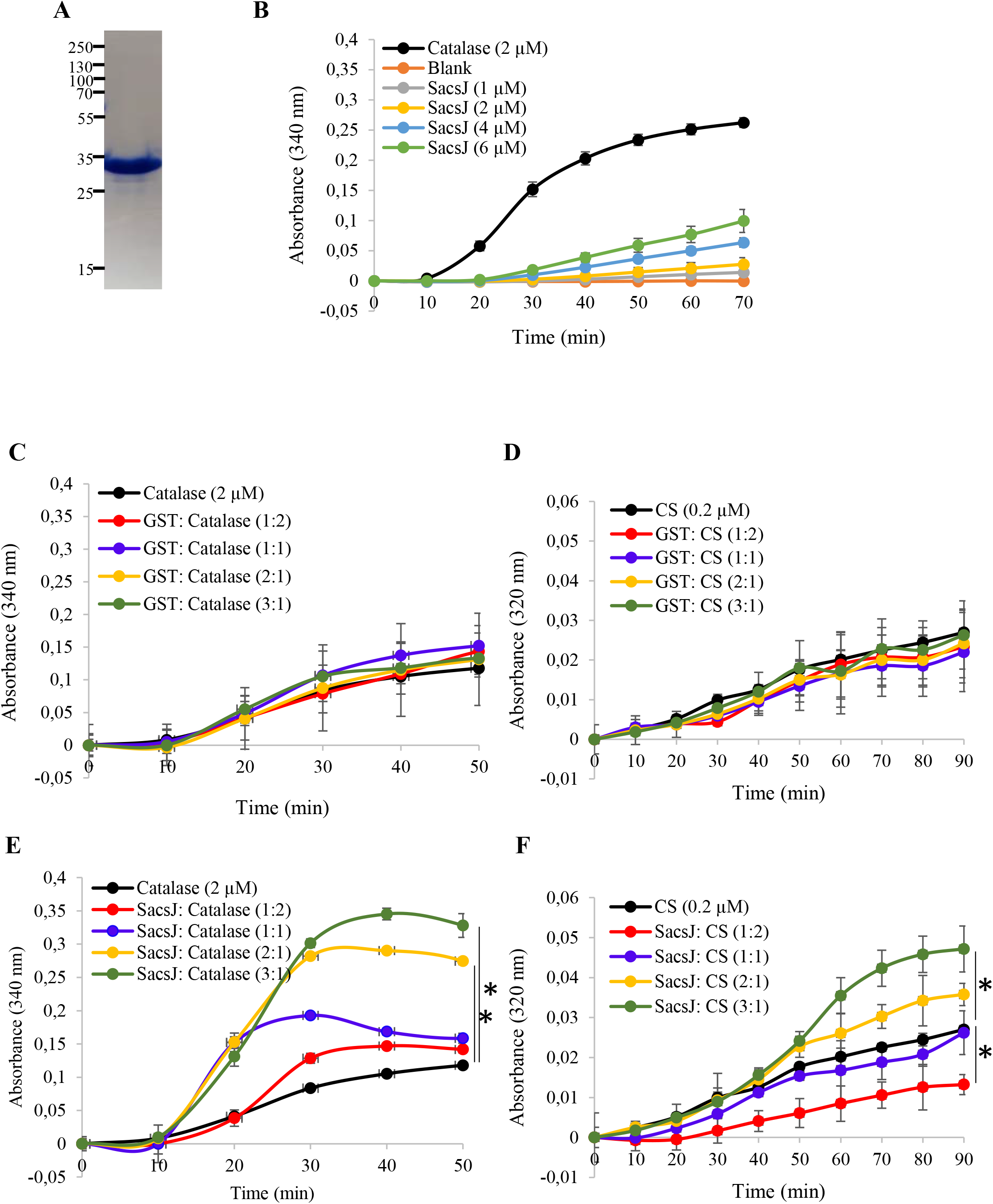
SacsJ domain did not act as a chaperone to reduce heat-denaturation of catalase or citrate synthase in an in vitro assay. A) Coomassie brilliant blue staining of a SDS-PAGE of purified recombinant SacsJ protein used in these assays. B) Absorbance at 340 nm, as a function of time in an assay for heat-denaturation of catalase (2 μM) at 60 °C compared to heat-induced denaturation in the presence of increasing concentrations of SacsJ-GST (1-6 μM). C-D, Absorbance at 340 nm of catalase (2 μM) or at 320nm citrate synthase (CS) (0.2 μM) incubated with or without GST (1-6 μM) heated at 60 °C for catalase or 45 °C for CS. (E-F) Absorbance at 340 nm and 320 nm of catalase (2 μM) or citrate synthase (CS) (0.2 μM) incubated with or without SacsJ-GST (1-6 μM) heated at 60 °C for catalase or 45 °C for CS. *p< 0.05 *vs* catalase or CS alone, one-way ANOVA, HSD Tuckey post hoc analysis (n=3).

### A cell-permeant SacsJ peptide disassembles vimentin filaments in fibroblasts

Our previous studies in cultured fibroblasts and motor neurons used plasmid delivery of myc-tagged sacsin domains [23], which is difficult to administer to neurons and other primary cells with low transfectability and challenging to apply therapeutically. We therefore developed a cell-permeant peptide (SacsJ-myc-TAT) in which the amino acid sequence of the SacsJ domain was fused to a myc epitope for detection and a cell penetrating sequence from the trans-activating transcriptional activator of HIV1 (TAT) for delivery into cells. GST-myc-TAT was also generated as a control. An immortalized line of the primary fibroblast strain MCH74 was used to establish dose-response for intracellular delivery of the peptide and its effect on vimentin IF. Fibroblasts were treated with 0.5 μM, 1 μM or 5 μM SacsJ-myc-TAT or GST-myc-TAT for 30 min, and the intracellular uptake of these peptides into the fibroblasts was detected by double label immunocytochemistry using antibodies against the myc-tag and vimentin and visualization of labeling by confocal microscopy. As shown in Fig. 4A, both peptides were detected in the cytoplasm and to a lesser extend in the nuclei of fibroblasts after 30 min, which is consistent with the biochemical properties of the TAT sequence [41, 42]. Fibroblasts receiving no treatment served as a control. Interestingly, the formation of perinuclear rings of vimentin (Fig. 4A) was a phenotype observed in fibroblasts treated with the SacsJ-myc-TAT for this short time period. At 0.5 μM the percentage of fibroblasts showing this phenotype was 38 ± 2.5% compared to 3.8 ± 1.3% in cultures treated with GST-myc-TAT and 2.3 ± 2.5 % with no treatment (Fig. 4B). This percentage was only slightly increased at higher concentrations of SacsJ-myc-TAT (1 or 5 μM) (Fig. 4B).

**Figure 4:**
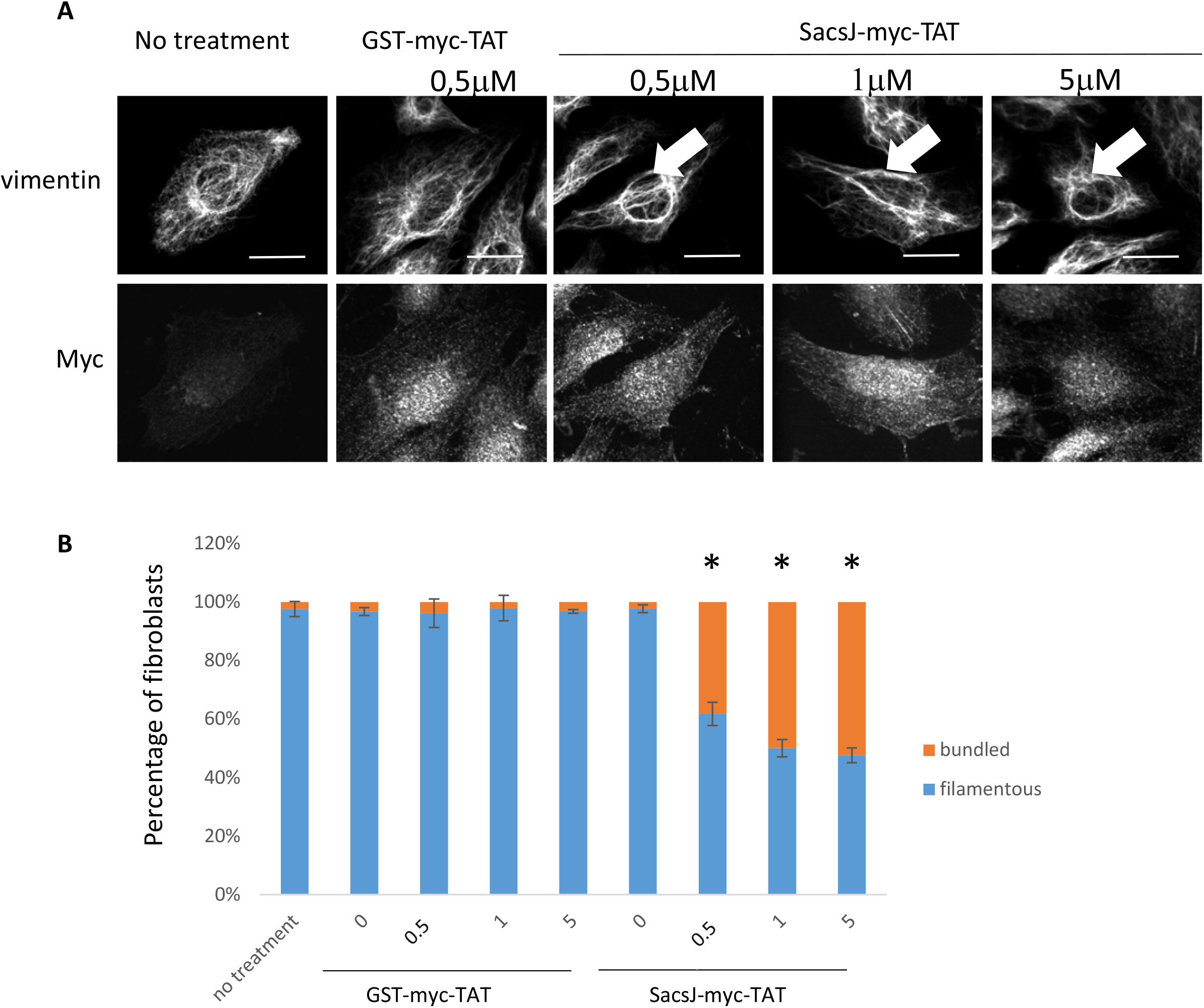
The cell-permeant peptide, Sacs-myc-TAT, induced step-wise disruption of the vimentin IF network in MCH74 fibroblasts. A) Representative 3-dimensional reconstructions of Z-stack confocal images of MCH74 fibroblasts double labelled with anti-myc (rabbit anti-myc) and anti-vimentin (V9 mouse monoclonal) treated with increasing concentrations of Sacs-myc-TAT as indicated (0.5 to 5 μM) or control peptide GST-myc-TAT. Treatment with Sacs-myc-TAT for 30 min resulted in nuclear rings of bundled vimentin IF (large arrow). Scale bar: 20 μm. B) Quantitation of the percentage of fibroblasts presenting circum-nuclear IF bundles or finely distributed IF when treated with increasing concentrations of Sacs-myc-TAT (0.5 to 5 μM) showing an increase in the percentage of cells with IF concentrated surrounding the nucleus. *p< 0.05 *vs* NFL alone, one-way ANOVA (n=3). Note the diffuse distribution of labelling of the myc tag on SacsJ.

Next, the effect of longer treatment durations with the SacsJ-myc-TAT or the GST-myc-TAT on the vimentin network were tested. Fibroblast cultures were treated with 0.5 μM SacsJ-myc-TAT or GST-myc-TAT for 0, 30 min, 3 h, 12 h or 24 h, followed by double immunolabeling for vimentin and the myc tag (Fig. 5). The GST-myc-TAT was detected intracellularly by indirect immunofluorescence up to 24 h suggesting a certain stability while, the SacsJ-myc-TAT was barely detectable at 24h. Qualitatively, the vimentin network appeared as a classic filamentous network in untreated fibroblasts or GST-myc-TAT-treated fibroblasts at all incubation times (Fig. 5A). On the other hand, fibroblasts treated with SacsJ-myc-TAT presented time-dependent phenotypes (illustrated in Fig. 5A and B). Perinuclear rings of bundled vimentin were observed at the 30 min time point as described above (Fig. 5B), but at 3 h, dismantling of the vimentin network had begun (Fig. 5C). By 12 h, cells showed diffuse, barely detectable, labelling of vimentin (Fig. 5D). At 24 h, a juxtanuclear focus of vimentin labelling was observed (Fig.5E), which together with a barely detectable SacsJ-myc-TAT suggests that this phenotype represents a recovery stage. As shown in Fig. 5F, the percentage of fibroblasts presenting perinuclear vimentin remained relatively constant during the first 12 h in the range of 36 ± 6.5 % to 44 ± 5.75 % (at 30 min and 12h of treatment, respectively) with no statistical difference by one-way ANOVA and a Tuckey HSD posthoc analysis. Similarly, the percentage of fibroblasts with a dismantled vimentin network remained constant from 3 h (20.64 ± 4 %) to 24 h (26.16 ± 1.43 %) while the percentage of fibroblasts with juxtanuclear vimentin (13.51± 1.3 %), which may represent recovery, increased. All together, our data suggest a dynamic regulation of the vimentin network by the SacsJ-myc-TAT leading to the stepwise disassembly of the vimentin network, which was captured by this experiment. Importantly, the SacsJ-myc-TAT peptide was taken up by the fibroblasts and reproduced the effect of the plasmid-derived ectopic expression of the SacsJ domain in disassembling the vimentin network [23].

**Figure 5:**
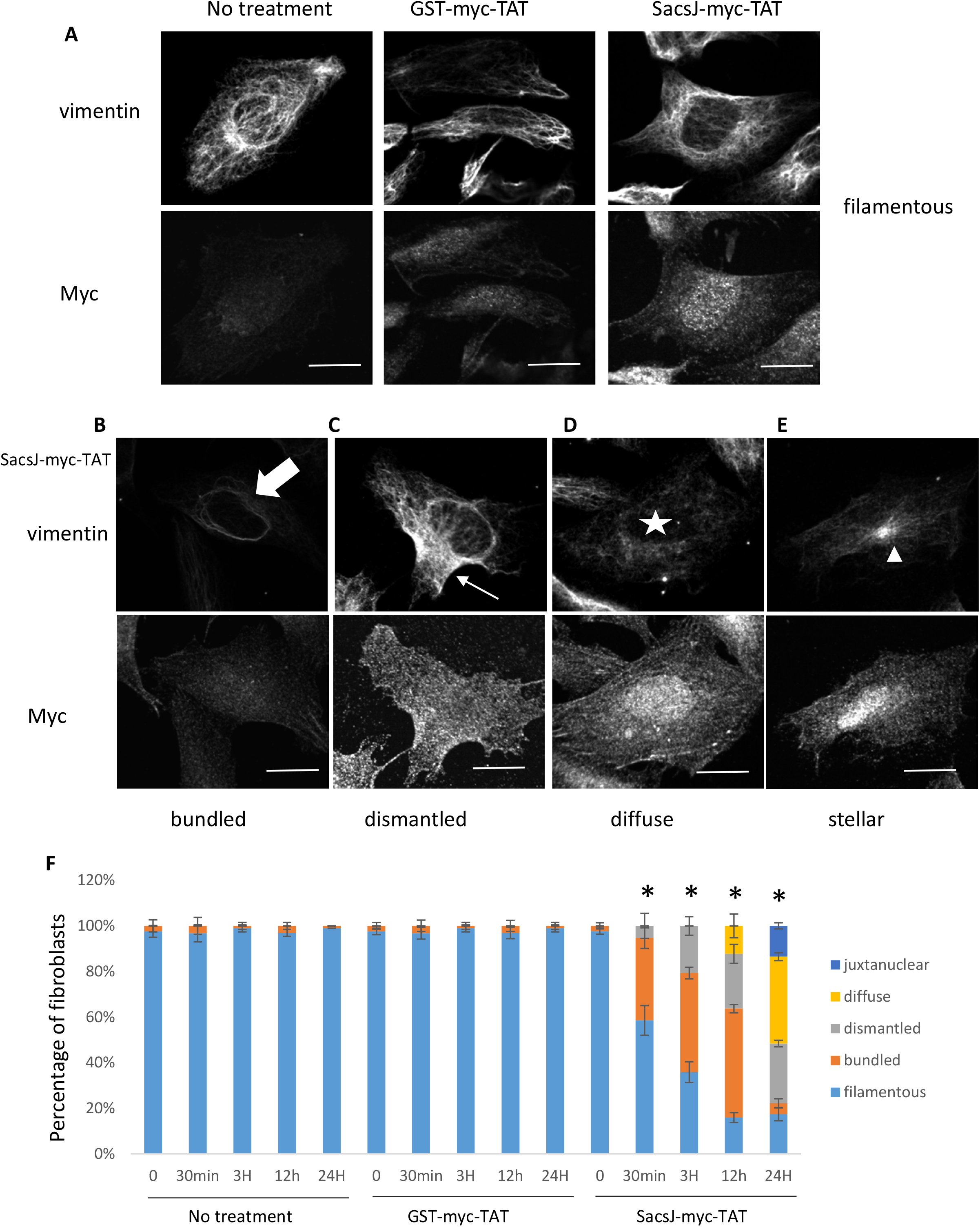
Over time, SacsJ-myc-TAT treatment resulted in the disassembly of vimentin IF in MCH74 fibroblasts. A) Representative 3-dimensional reconstructions of Z-stack confocal images of fibroblasts double labelled with anti-myc (rabbit anti-myc) and anti-vimentin (V9 mouse monoclonal). Scale bar: 20 μm. (B-E) MCH74 fibroblasts were treated with GST-myc-TAT control or SacsJ-myc-TAT (0.5 μM) for 30 min, 3 h, 12 h or 24 h and showed time-dependent phenotypes: perinuclear rings of bundled vimentin (B, large arrow), dismantled vimentin network (C, small arrow), diffuse vimentin labelling (D, star), and appearance at a juxtanuclear accumulation (E, arrowhead). F) Quantitation of the percentage of fibroblasts presenting those phenotypes over duration of SacsJ-myc-TAT treatment. *p< 0.05 *vs* time-matched no treatment or treated with GST-myc-TAT (0,5μM) one-way ANOVA, HSD Tuckey post hoc analysis (n=3).

### SacsJ disassembles NF in mouse motor neurons in culture

To determine the efficiency of SacsJ-Myc-TAT to resolve the NF network and bundles in sacsin wild-type and sascin-deficient neurons, motor neurons in 6 week-old *Sacs*^+/+^ or *Sacs*^-/-^ spinal cord-DRG cultures were treated with 0.5 μM SacsJ-Myc-TAT or its control, GST-myc-TAT, for 30 min. Intracellular uptake of SacsJ-Myc-TAT or GST-myc-TAT was detected by anti-myc immunolabeling and the NF network was visualized and evaluated by double-labelling cells with anti-NFL (Fig. 6A and Fig.7A). Qualitatively, the NF network in *Sacs*^+/+^ motor neurons either remained filamentous or was dismantled/disassembled (Fig. 6A and B) in cultures treated with the SacsJ-Myc-TAT peptide. In 6 week-old *Sacs*^-/-^ spinal cord-DRG cultures, 83 ± 5% of motor neurons contained well-established NF bundles (Fig. 7A), which decreased to 22.08 ± 5% of neurons following treatment with SacsJ-myc-TA, with GST-myc-TAT having no significant effect (Fig. 7B). Qualitatively, the NF networks in *Sacs*^-/-^ motor neurons treated with GST-myc-TAT were more spatially distributed rather than bundled in dendrites and cell bodies.

**Figure 6:**
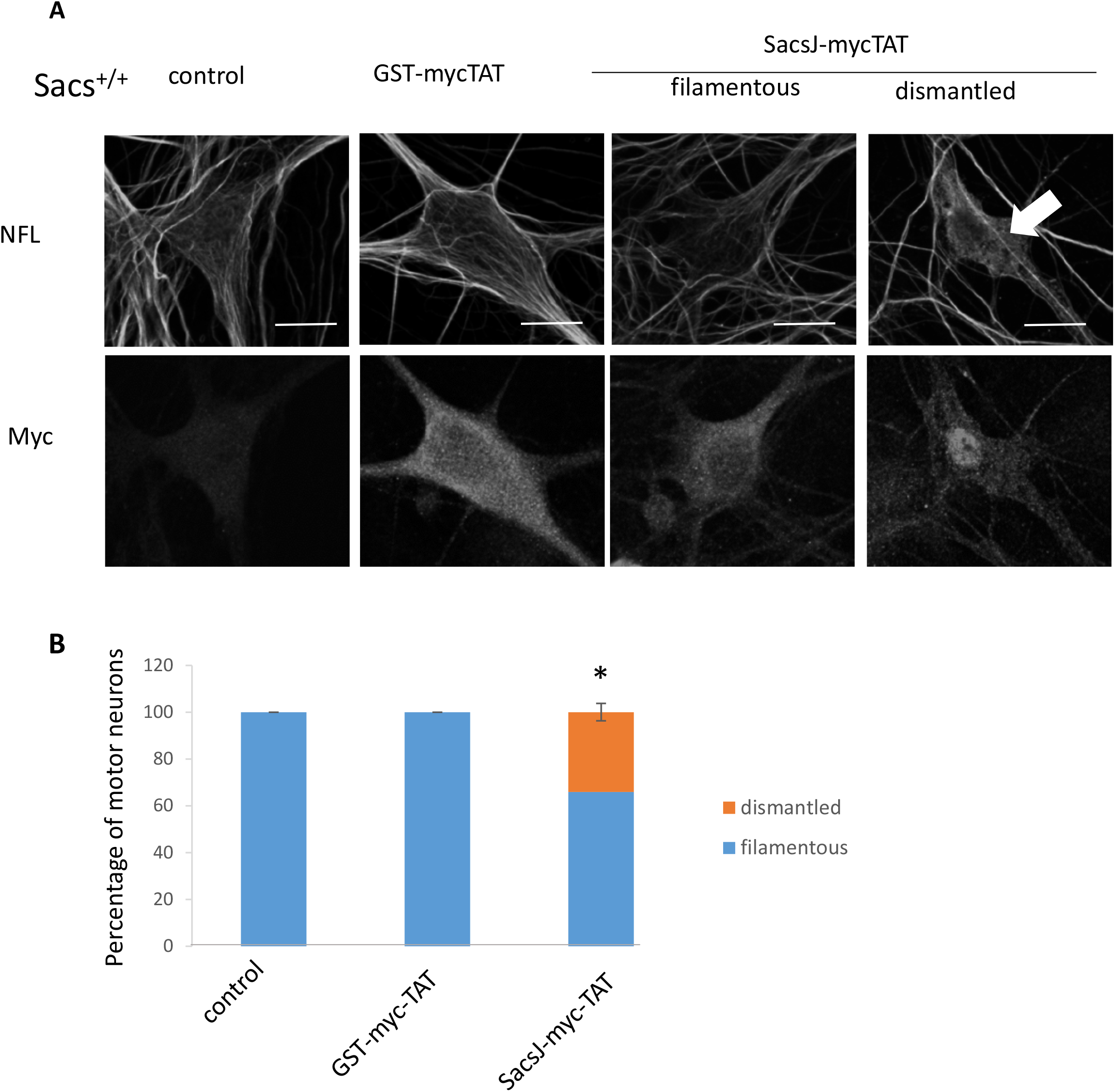
The cell-permeant peptide, SacsJ-myc-TAT, induced the disassembly of NF in *Sacs*^+/+^motor neuron in culture. A) Representative 3-dimensional reconstructions of Z-stack confocal images of double labelling with anti-myc (rabbit anti-myc) and anti-NFL in *Sacs*^+/+^ 6 week-old murine spinal cord-DRG cultures showing the NF network and distribution of myc-TAT peptides in motor neurons. Cultures were treated with SacsJ-myc-TAT (0.5 μM) or GST-myc-TAT control peptide for 30 min and compared to untreated cultures. SacsJ-myc-TAT dismantled the endogenous NF network (large arrow). Scale bar: 20 μm. (B) Quantitation of the percentage of motor neurons presenting a filamentous or dismantled NF network when treated with Sacs-myc-TAT (0.5 μM). *p< 0.05 *vs* no treatment or treated with GST-myc-TAT (0.5μM) using a one-way ANOVA, HSD Tuckey post hoc analysis (n=3).

**Figure 7:**
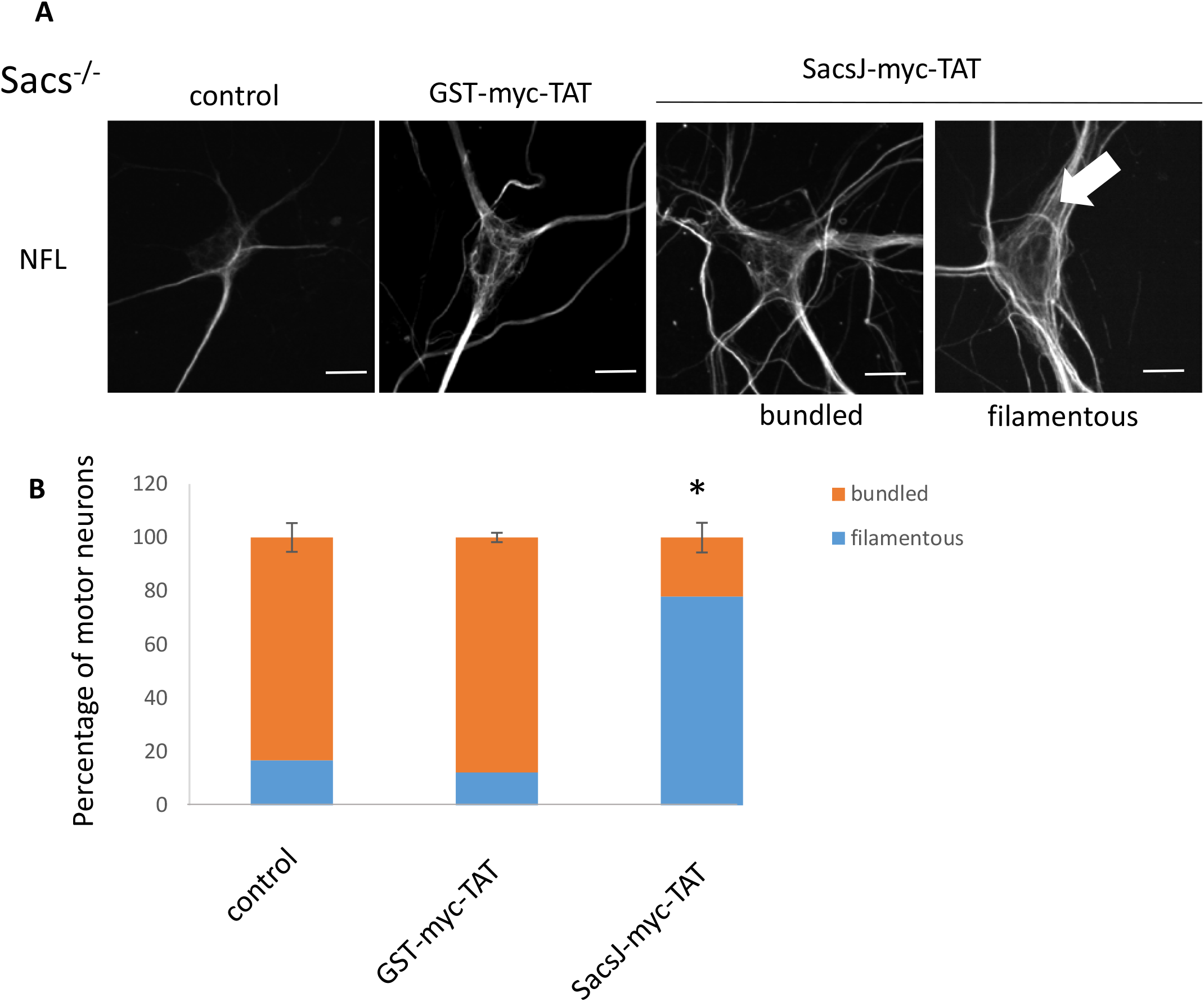
The cell-penetrating peptide, SacsJ-myc-TAT, resolved NFL bundles in *Sacs*^-/-^ motor neuron in culture. A) Representative 3-dimensional reconstructions of Z-stack confocal images of motor neurons in *Sacs*^-/-^ 6 week-old spinal cord-DRG cultures double labelled with anti-myc (rabbit anti-myc) and anti-NFL to show the NF network and SacsJ-myc-TAT distribution in motor neurons. Cultures and were treated with SacsJ-myc-TAT (0.5 μM) or GST-myc-TAT control peptide for 30 min and compared to untreated cultures. Treatment with SacsJ-myc-TAT resolved the NF bundles (large arrow). Scale bar: 20 μm. (B) Quantitation of the percentage of motor neurons presenting a filamentous or bundled NF network when treated with Sacs-myc-TAT (0.5μM). *p< 0.05 *vs* no treatment or treated with GST-myc-TAT (0.5μM) using a one-way ANOVA, HSD Tuckey post hoc analysis (n=3).

## Discussion

This study demonstrated a direct role of the SacsJ domain of sacsin in IF assembly/disassembly. In vitro, SacsJ disrupted both pre-existing NFL filaments and the formation of new NFL filaments, in the absence of any Hsp70. There was no evidence of general chaperone activity of SacsJ, suggesting that its effects are specific to NFL, or more broadly to IF subunits. The cell penetrating peptide SacsJ-myc-TAT caused disruption of the vimentin network in fibroblasts. It efficiently cleared the abnormal bundles of NF in *Sacs*^-/-^ motor neurons, but also caused partial dismantling/disassembly of the NF network in *Sacs*^+/+^ neurons. These results suggest that SacsJ itself is a potent regulator of NF and IF networks.

While assembly of IF proteins is mainly regulated by phosphorylation of N-terminal residues [28], some small heat shock protein chaperones, HSPB1 and HSPB6, can prevent the assembly of IF, disassemble IF into dimers and tetramers, and maintain IF proteins soluble in non-denaturing detergent [29, 31, 43]. In contrast to HSPB1 or HSPB6, the SacsJ domain did not solubilize or co-sediment with NFL in vitro despite the fact that NFL filaments were dismantled or prevented from assembling. When incubated with the SacsJ domain, NFL pelleted following centrifugation, rather than being retained in the supernatant as would dimers and tetramers [31], but the absence of intact filaments and shorter filaments corresponding to UFL by TEM indicates the presence of some intermediate sized form of NFL in addition to aggregates in the presence of SacsJ.

DNAJ is a large family of proteins beneficial in restoring neuronal proteostasis and reducing neurotoxicity associated with proteins aggregation. The specificity of DNAJ toward certain client-proteins envisions individual DNAJ proteins to be tailored to distinct aggregation-prone proteins [44]. A few DNAJ proteins have been shown to play a role in IF assembly and homeostasis. For example, DNAJB6/MrJ, which interacts with keratin K8/K18, is a cochaperone recruiting HSC70 to organise the keratin IF network; intracellular microinjection of a neutralising antibody to DNAJB6 resulted in abnormal bundling of keratin filaments [34]. In fact, HSP40 has been shown to regulate keratin proteins via the ubiquitin-proteasome pathway and is believed to play a role in the exchange of the Keratin5/Keratin14 pair to the Keratin1/Keratin10 pair into the keratin filaments during maturation of HaCaT keratinocytes. This regulation occurs through degradation of Keratin1 [45]. The J domain of sacsin could play a similar role in regulation of IF protein dynamics.

We also developed a cell-penetrating peptide SacsJ-myc-TAT that retained SacsJ properties identified previously by plasmid-mediated delivery [23], specifically, disassembly of the vimentin IF network in fibroblasts. The peptide delivery system facilitated examination of the effects of SacsJ on the network over time in vivo and revealed a stepwise disorganisation of vimentin IF, starting by depletion of IF in the periphery and concentrating around the nucleus and progressing to loss of bundles and overall vimentin labeling. The bundles around the nucleus are reminiscent of the perinuclear rings of bundled vimentin, described in endothelial cells or lymphoblasts undergoing mitosis or when treated with colcemid [46–48] or PP2A inhibitor [49]; i.e., they are first to form and last to go. Indeed, such rings of bundled vimentin appear within the first hours of cell adhesion [18, 50] and may reflect a difference of IF dynamics between the periphery and the perikaryon. Indeed, we showed that NF organized in bundles in *Sacs*^-/-^ motor neurons in culture had a slower turn over of subunits than normal NF network [23].

The cell-penetrating peptide SacsJ-myc-TAT was also tested in spinal cord-DRG cultures, demonstrating dissolution of NF in motor neurons. In cultures prepared from *Sacs*^-/-^ mice, SacsJ-myc-TAT resolved the characteristic NF bundles. Whereas a more normal NF network was preserved in *Sacs*^-/-^ neurons treated with the SacsJ peptide, the loss of IF networks in *Sacs*^+/+^ fibroblasts or neurons indicate that use of the SacsJ domain alone therapeutically could be problematic over the longer term or dosage will be crucial. In contrast, overexpression or induction of the HSP70/HSPA1 resolved bundles in *Sacs*^-/-^ motor neurons in culture without destroying the normal network. Nonetheless, abnormally high levels of the general chaperone HSP70/HSPA1 may have other unwanted effects compared to the expression of smaller domains with specific functions. The challenge with sacsin replacement therapy is that the sequence is too large for delivery methods. We are currently examining other sacsin domains and domain combinations with SacsJ in order to design an effective miniconstruct.

## ACKNOWLEDGEMENTS

We thank Jeannie Mui and Dr Vali at the Facility for Electron Microscopy Research of McGill University for help in microscope operation and data collection. This study was supported by grants from the Ataxia of Charlevoix-Saguenay Foundation to B. J. Gentil and H.D. Durham and Muscular Dystrophy Canada. The authors declare no competing financial interests.

## Author contributions

A. Dabbaghizadeh performed the in vitro experiments and analyzed the majority of the data. B.J. Gentil and H.D. Durham conceived, designed, supervised the project and wrote the manuscript, with key input from the other authors. A. Paré, Z. Cheng-Boivin and R. Dagher carried out the cellular experiments. S. Minotti produced and maintained spinal cord cultures. B. Brais and MJ Dicaire provided the *Sacs*^-/-^ mice and reviewed the manuscript. J. Young provided key insights and suggestions for the experiments as well as reviewed the manuscript.

## Abbreviations

ARSACS: Autosomal Recessive Spastic Ataxia of the Charlevoix Saguenay
IF: intermediate filaments
NF: neurofilaments
TEM: transmission electron microscopy
CS: citrate synthase
ULF: Unit-Length Filaments
HIV1: Human Immunodeficiency Virus1
Ubl: ubiquitin-like domain
NFL: neurofilament light polypeptide
SacsJ: Sacsin J domain
SIRPT: Sacsin Internal RePeaT domain
HEPN: Higher Eukaryote and Prokaryote Nucleotide-binding domain

